## Supplementary Material for "Endothelin 3 and T-type Ca^2+^ channels drive enteric neural crest cell calcium activity, contractility and migration"

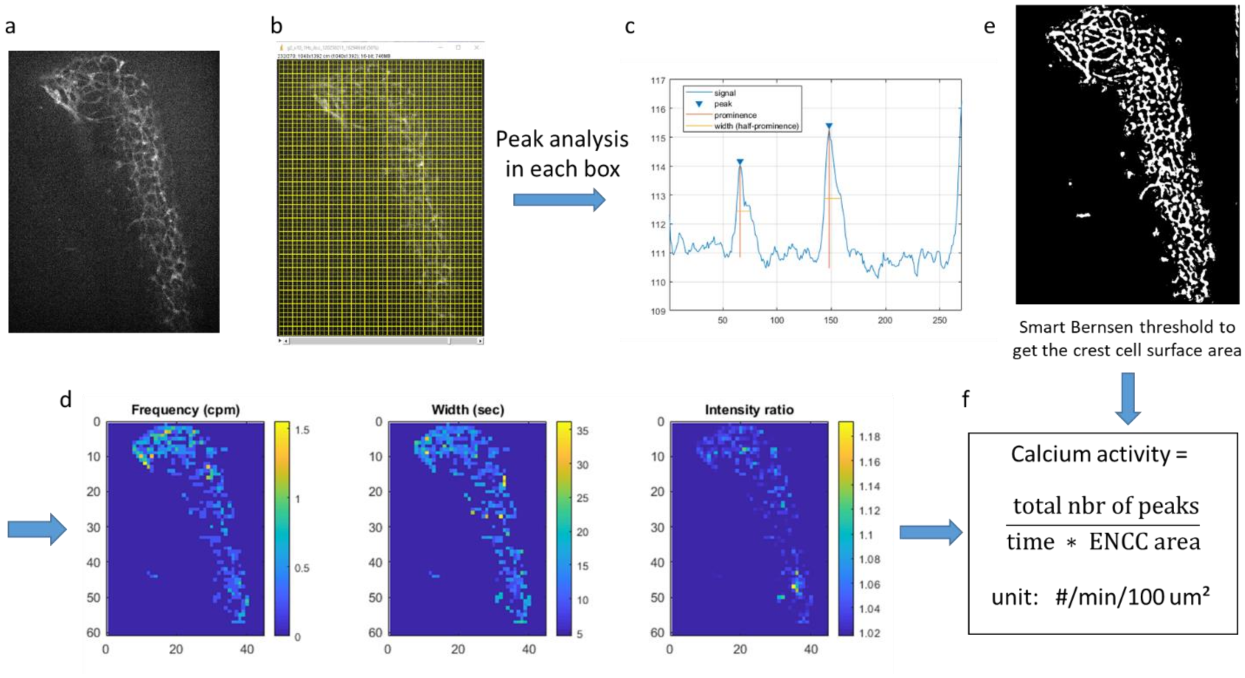


***Figure S1. Methodology of $Ca^{2+}$ activity analysis.*** *(a) The 8-bit video acquired at x10 magnification is loaded in ImageJ. (b) The Macro "gridROI" is applied to overlay a grid of 45x61=2745 square boxes, each square has an area of 250 $\mu m^2$. The "Multimeasure" command from the ROI Manager measures the grayscale intensity in each square over time. (c) This data is input in Matlab for automatic peak detection. The thresholds used for all analysis are: minimum peak prominence 2, minimum peak width 3 sec, minimum peak-to-peak distance 3 sec. (d) Heatmaps of frequency (F), width (W) and intensity ratio (IR) are output, as well as the average of these values over the whole sample, and the total number of peaks detected. (e) To compute the total area occupied by ENCCs, the time-averaged intensity of the video is first projected, the resulting imaged Gaussian blurred (radius 3), and a Bernsen local threshold is applied, with radius 15 px, starting at parameter1=12. The parameter1 is gradually decreased by steps of 1, including more and more material within the threshold. At each step, the difference between the previous and new average gray intensity is computed; when this difference exceeds a certain constant threshold (2), the loop is ended and the last parameter1 value is saved and used for calculation of the threshold and of the resulting ENCC area. This approach enables to perform the threshold automatically, in a user-independent way. (f) The calcium activity is computed by dividing the total number of events by the time of the recording (in min) and by the total area of the ENCCs (in units of one cell area, ~100 um²). The whole analysis procedure has been automated for batch processing of time-lapse stacks.*

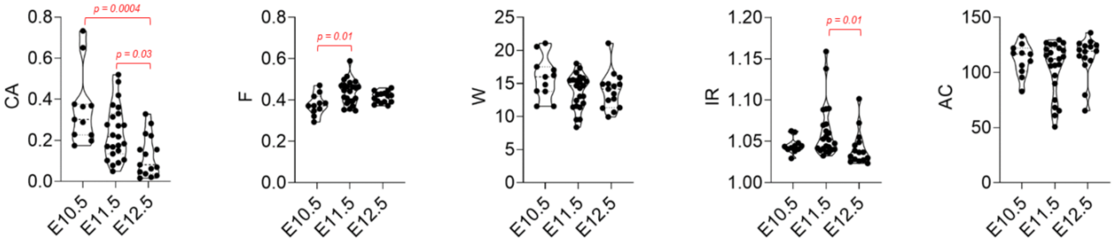


***Figure S2. CA characteristics across stages at the migration front.*** *CA = calcium activity in events/min/100 µm², F = frequency in cycles-per-min, W=width in sec, IR= intensity ratio (dimensionless), AC=average calcium in arbitrary pixel unit. Only statistically significant differences (p<0.05) are indicated, Kruskall Wallis test.*

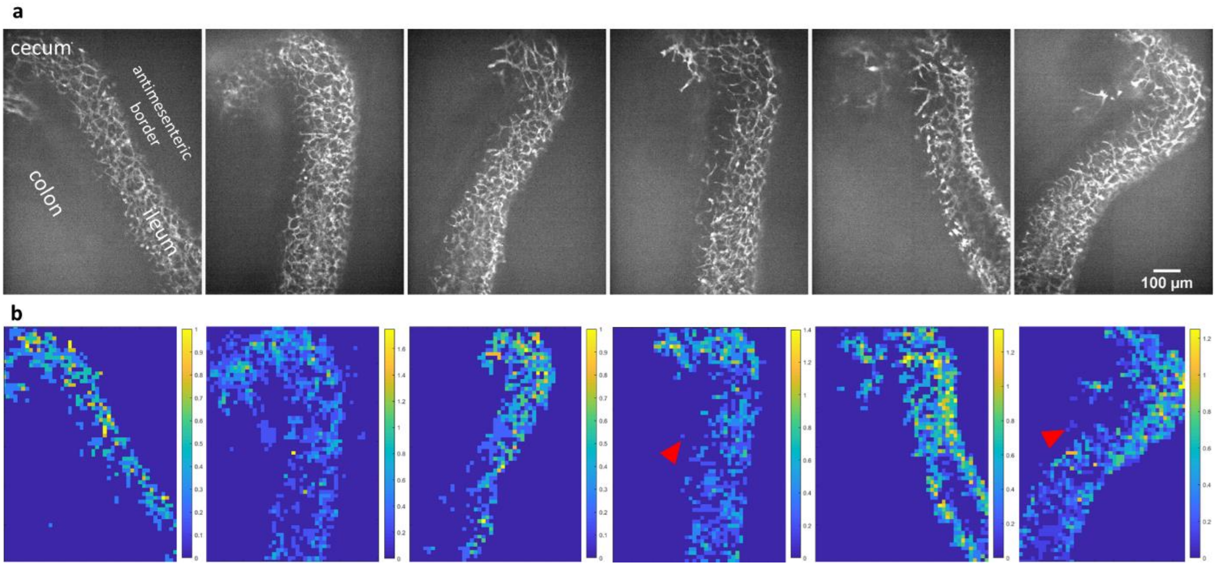


***Figure S3. Calcium transients are concentrated at the cecum and ileum anti-mesenteric border.*** *(a) 6 different E11.5 samples at the ileo-cecal junction, time-averaged projection of 3-4 min long 1Hz timelapse. The cecum is a the top, the ileum at the bottom, and the antimesenteric border to the right, the uncolonized colon (not visible) to the left. (b) Corresponding frequency heatmaps (scalebar unit: cpm). $Ca^{2+}$ activity is most important at the cecum and antimesenteric ileum border. Red arrowheads point to activity in transmesenteric ENCCs.*

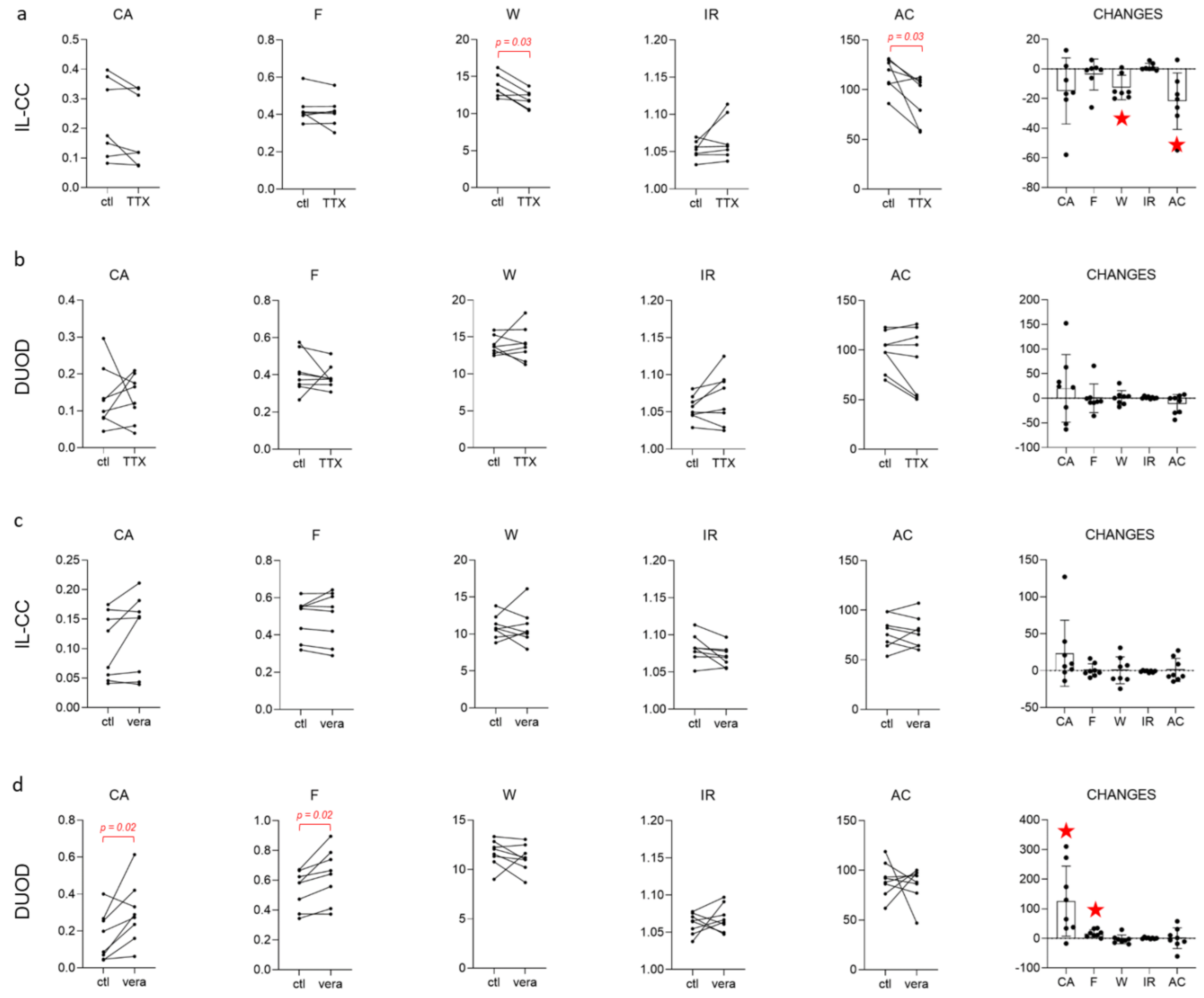


***Figure S4. Role of $Na^+$ channels in $Ca^{2+}$ activity.*** *Effects of tetrodotoxin 1 µM at E11.5, at the ileo-cecal junction (a) and in the duodenum (b), and of veratridine 5 µM at E11.5, at the ileo-cecal junction (c) and in the duodenum (d). CHANGES are* $100 * (x_{drug} - x_{control})/x_{control}$ *in all figures of the manuscript. Only statistically significant differences (p<0.05 and red stars) are indicated, unless otherwise stated the Wilcoxon matched-pairs signed rank test was used in the Supplementary Material part.*

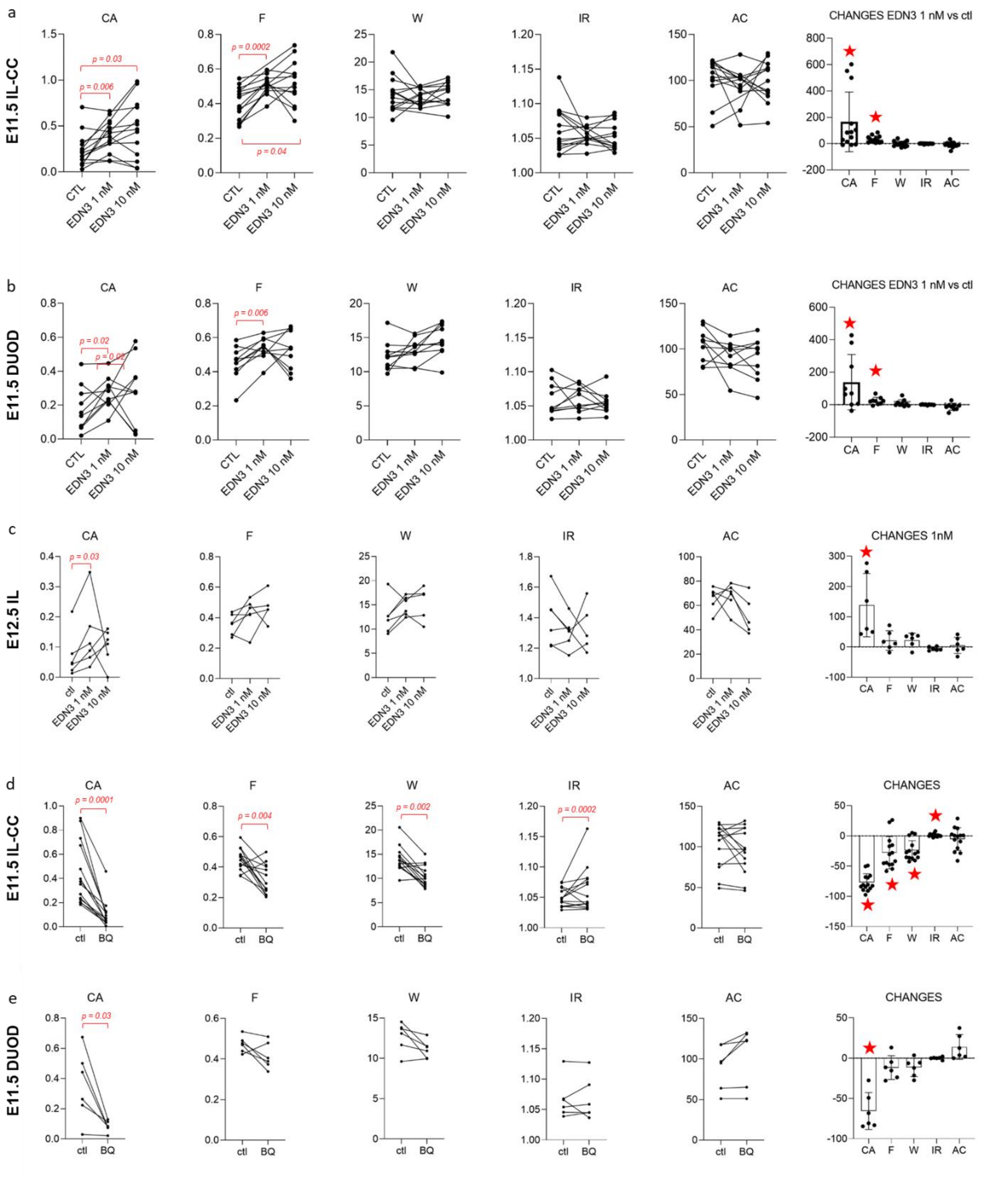


***Figure S5. Endothelin 3 and receptor EDNRB are critical for $Ca^{2+}$ activity**. (a,b) Effects of EDN3 1 nM and 10 nM at E11.5, at the ileo-cecal junction (a) and in the duodenum (b). (c) Effects of EDN3 1 nM and 10 nM at E12.5, in the ileum. n=5/6 in this assay exhibited a $Ca^{2+}$ surge in all ENCCs upon application of EDN3 10 NM. In n=1/5 samples that exhibited a $Ca^{2+}$ surge, CA was reduced post-flash, and absent in n=1/5. In the remaining samples CA was increased compared to the control situation. (d,e) Effects of receptor EDNRB blockade by BQ 788 10 µM at E11.5, at the ileo-cecal junction (c) and in the duodenum (e).*

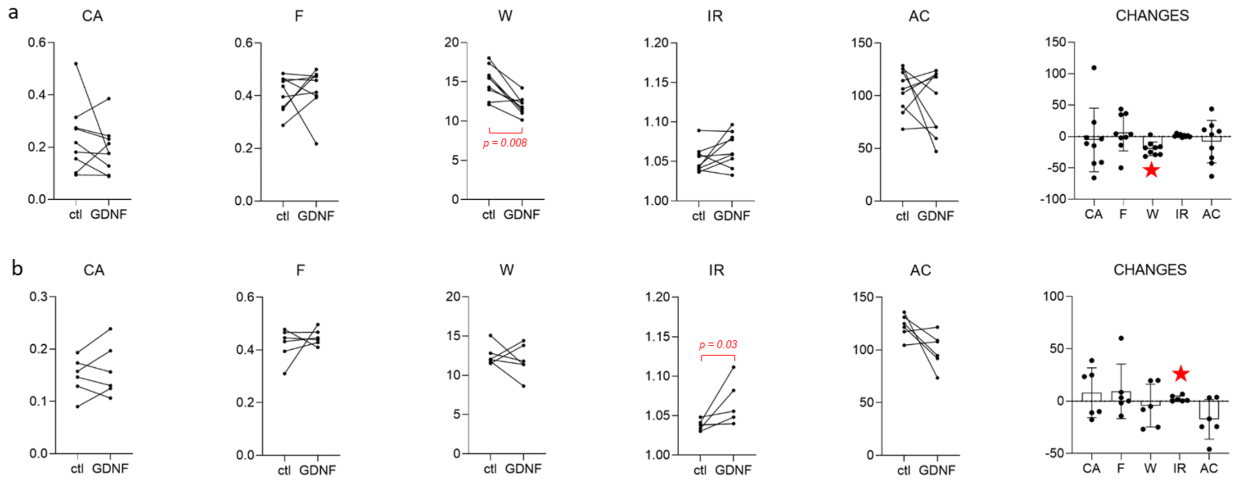


***Figure S6. Effects of GDNF 10 ng/mL at E11.5.*** *(a) At the ileo-cecal junction. (b) In the duodenum.*

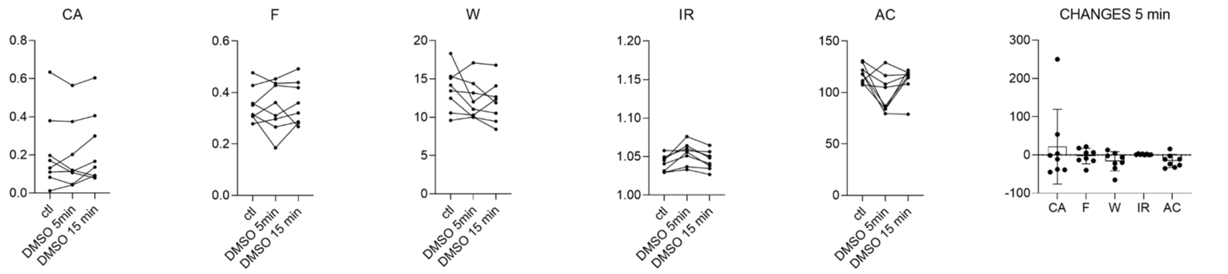


***Figure S7. Effect of DMSO vehicle on $Ca^{2+}$ activity.*** *A volume ratio of 2 µL DMSO / 1000 µL medium was applied, corresponding to the maximum ratio used in this report. Calcium imaging was performed 5 min and 10-20 min post-application. None of the differences were significant.*

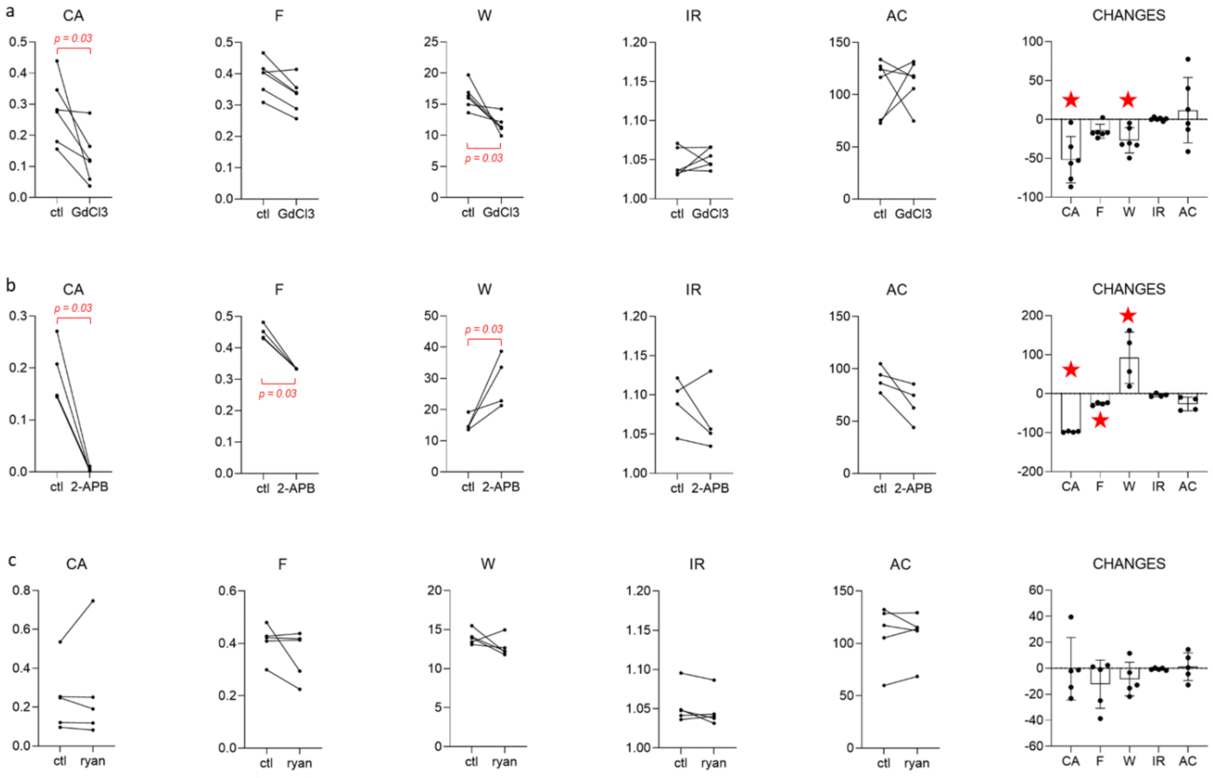


***Figure S8. Extracellular and intracellular sources of $Ca^{2+}$.*** *Effects of $GdCl_3$ 100 µM (a), 2-APB 100 µM (b), ryanodine 10 µM (c) at E11.5, at the ileo-cecal junction. p-values in (b) are obtained from the Mann-Whitney two-tailed test.*

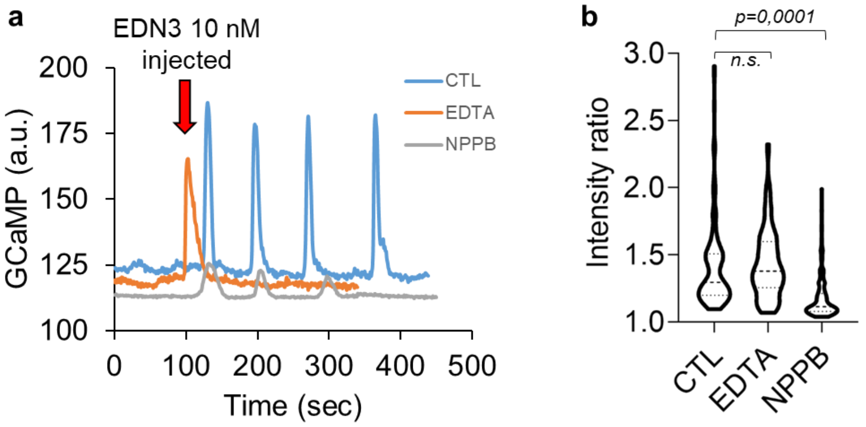


***Figure S9. Induction of CA by EDN3 10 nM in control, EDTA and NPPB conditions.*** *(a) A few $Ca^{2+}$ transients could still be elicited by administration of EDN3 10 nM after application of EDTA or of NPPB. (b) Transient intensity ratios measured from ROIs surrounding individual cells after EDN3 10 nM administration for control (160 cells analyzed from n=6 samples), EDTA (120 cells, n=6) and NPPB conditions (230 cells, n=6).*

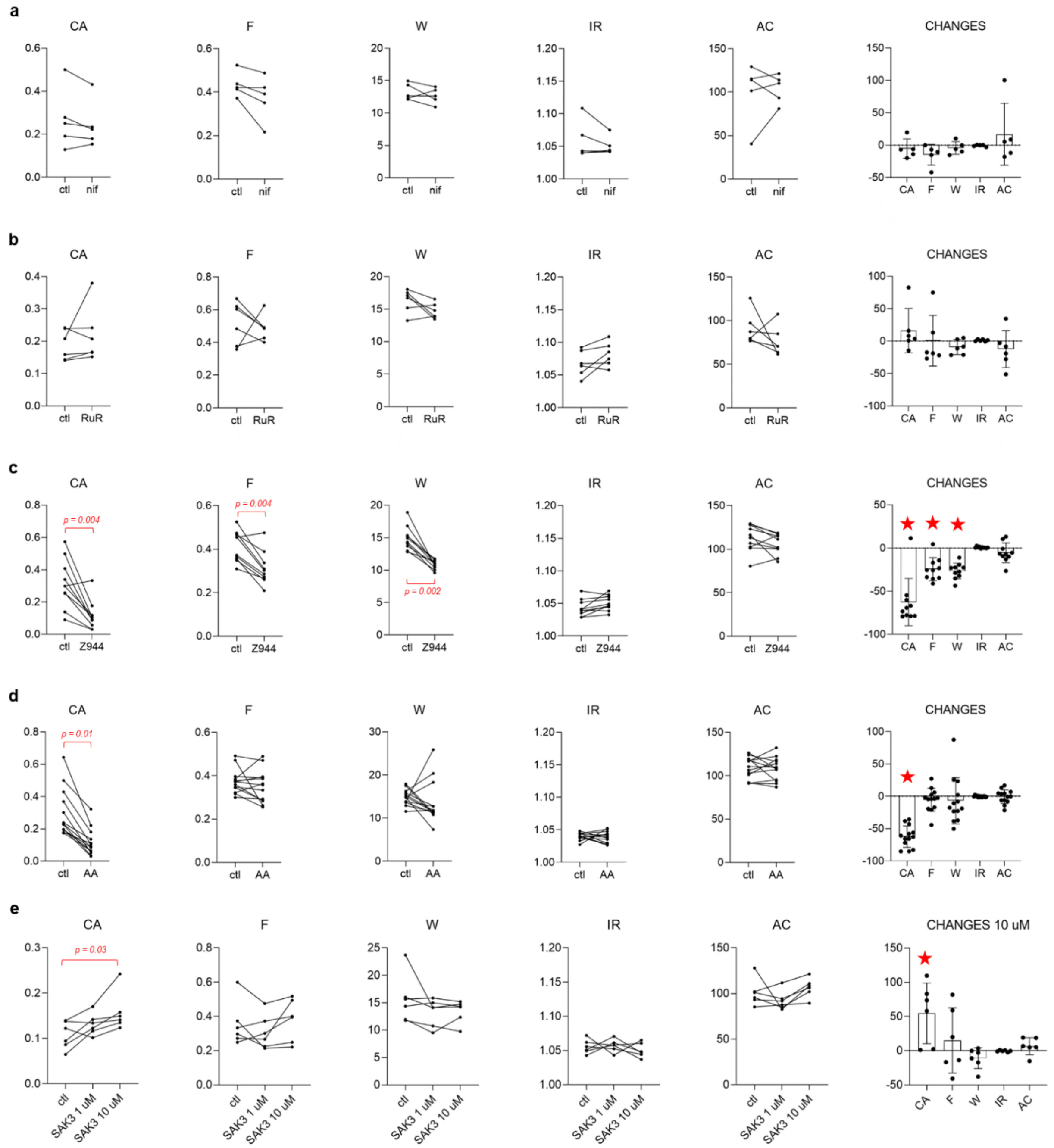


***Figure S10. $Ca^{2+}$ channel dependence of CA.*** *Effects of nifedipine 10 µM (a), ruthenium red 100 µM (b), Z944 50 µM (c), ascorbic acid 1 mM (d) and SAK3 1 & 10 µM (e) at E11.5, at the ileo-cecal junction.*

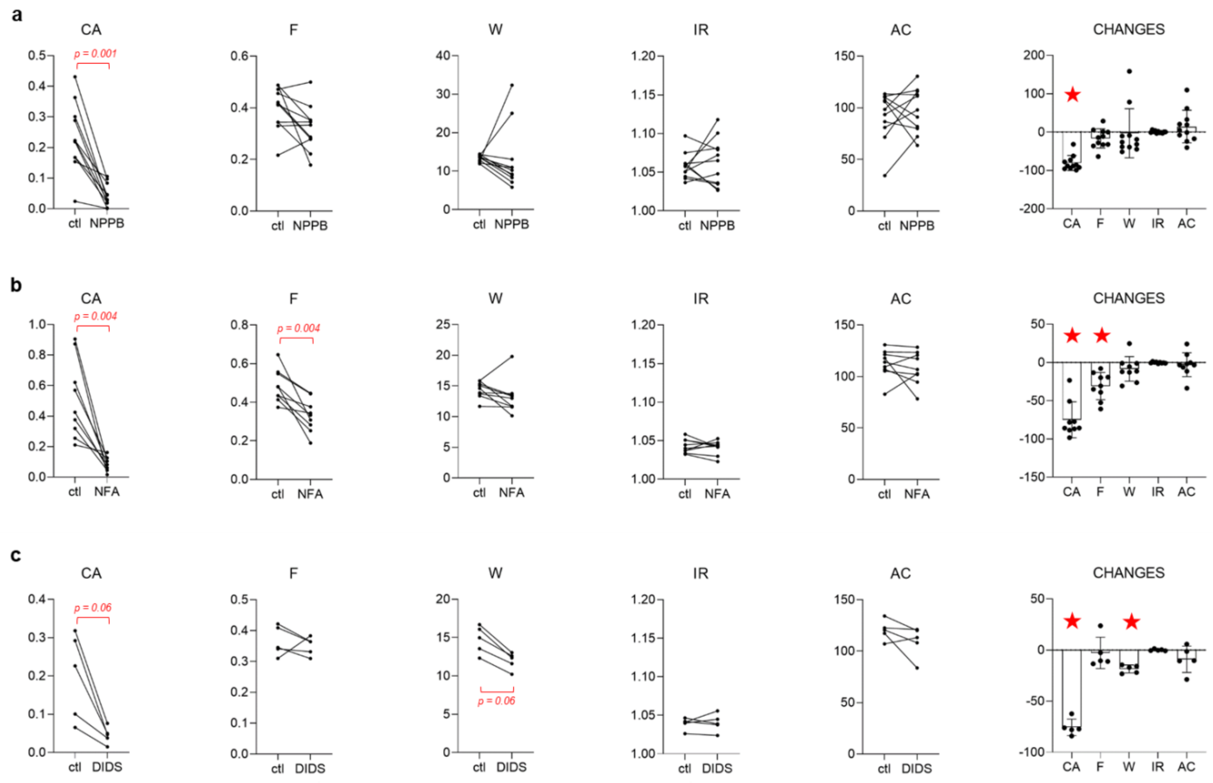


***Figure S11. Cl- channel dependence of CA.*** *Effects of NPPB 100 µM (a), NFA 50 µM (b) and DIDS 500 µM (c) at E11.5, at the ileo-cecal junction.*

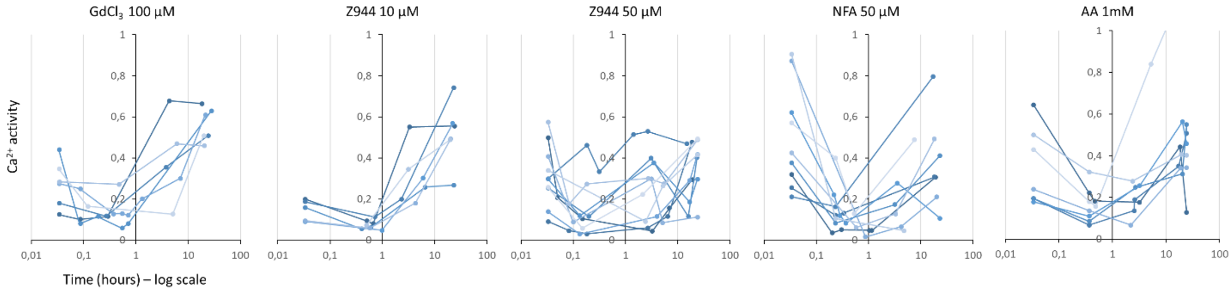


***Figure S12. Kinetics of $Ca^{2+}$ activity inhibitors.*** *The time is indicated in log-scale for clarity: points on the very left are before drug application, CA is followed up to 24 hours. CA increases sharply after 1h for $GdCl_3$ 100 µM and Z944 10 µM above initial levels. $GdCl_3$ was unstable in the culture medium, precipitating out after a few hours (visible as a fine powder that gathered towards the dish center when the Petri was gently swirled), which likely explains the kinetics for this drug. We noticed that the effectiveness of Z944 strongly decreased following reconstitution in DMSO and freeze-thaw cycles: it is therefore likely that the time-dependent behavior is due to degradation of the drug over the course of the experiment. For Z944 50 µM, NFA 50 µM and AA 1mM, the reduction of CA lasts ~10h, and the CA at 24h was on the same level as it was initially.*

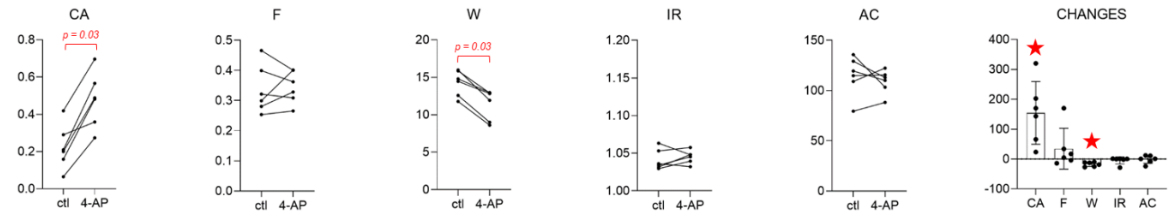


***Figure S13. K⁺ channel dependence of CA.*** *Effect of 4-AP 1 mM at E11.5, at the ileo-cecal junction.*

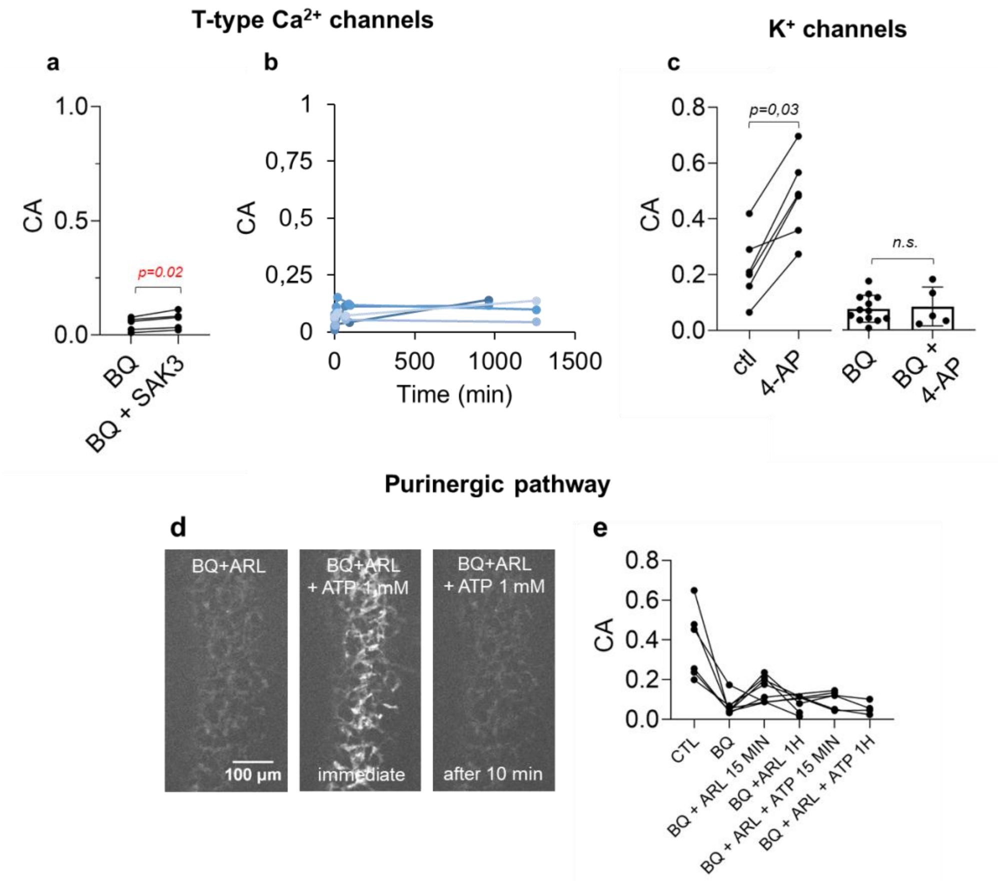


***Figure S14. Stimulation of T-type $Ca^{2+}$ channels, $K^{+}$ channels or purinergic receptors after EDNRB blockade does not allow to recover physiological CA levels****. (a) CA of BQ788 10 µM treated E11.5 ilcc samples, and 10 min after application of SAK3 10 µM, n=5, paired Student t-test. The increase is significant, but the CA remains far below the one measured in the absence of BQ (Fig.2e). (b) CA activity of n=5 BQ788 10 µM + SAK3 10 µM remains at a low level throughout the 1 day culture period. (c) Left: effect of 4-AP 1 mM on CA on control guts (WMP test). Right: CA comparison of BQ788 (n=14) and BQ788 + 4-AP 1 mM treated samples at E11.5 (n=5, Mann-Whitney test). (d) $ATP_e$ could elicit a strong $Ca^{2+}$ rise in all ENCCs in the field of view upon acute administration, but CA returned to low levels after this (Video S8). (e) Follow-up of n=8 samples first treated with BQ and then stimulated via the purinergic pathway, with ARL 67156 alone (100 µM) and with further addition of $ATP_e$ (100 µM). CA remained at low levels.*

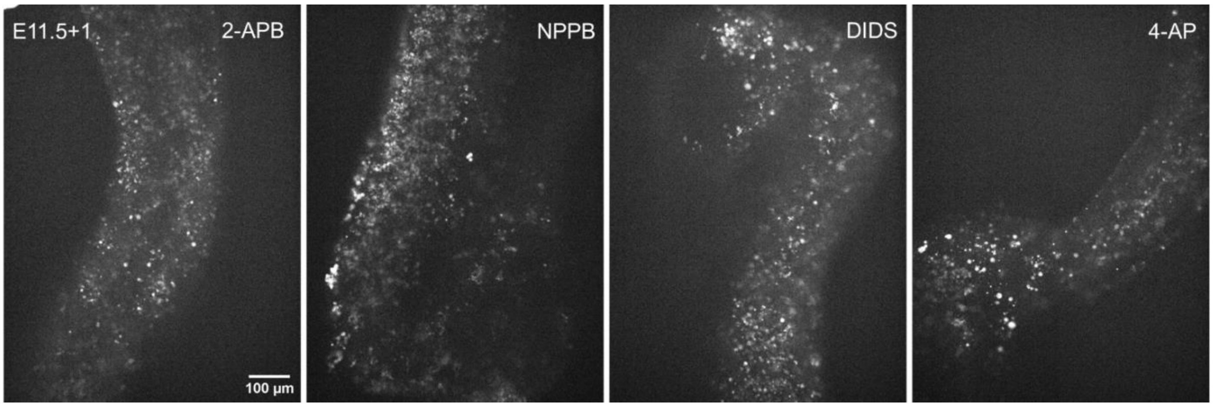


***Figure S15. 2-APB 100 µM, NPPB 100 µM, DIDS 500 µM, 4-AP 1 mM induced massive ENCC death after 24h application.*** *Cells are disconnected, round, small, with high intracellular $Ca^{2+}$. These patterns were systematic in all samples examined*

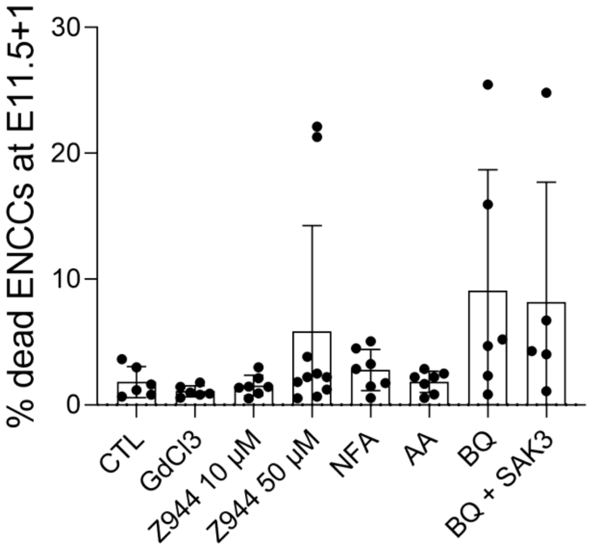


***Figure S16. Quantification of cell death in E11.5+1 guts in different culture conditions.*** *x10 z-stqck imqges of the samples after 1 day culture at the ilcc junction were max z-projected (as in Fig.5a). In ENCC-specific GCaMP samples, dead cells appear as round and brilliant because they are loaded with $Ca^{2+}$ (the membrane is disrupted). The percentage of dead ENCCs was computed from the ratio of the surface occupied by these round brilliant cells versus the surface occupied by all ENCCs. The fraction of dead cells was <7% in 49/54 samples, and none of the fractions were significantly different (Mann-Whitney test and Student t-test) from the control group, although BQ788 tended to induce higher average cell death. High cell death (10-25 %) was found in n=2/10 Z944 50 µM samples, in n=2/6 BQ 10 µM samples and in n=1/5 BQ + SAK3 samples.*

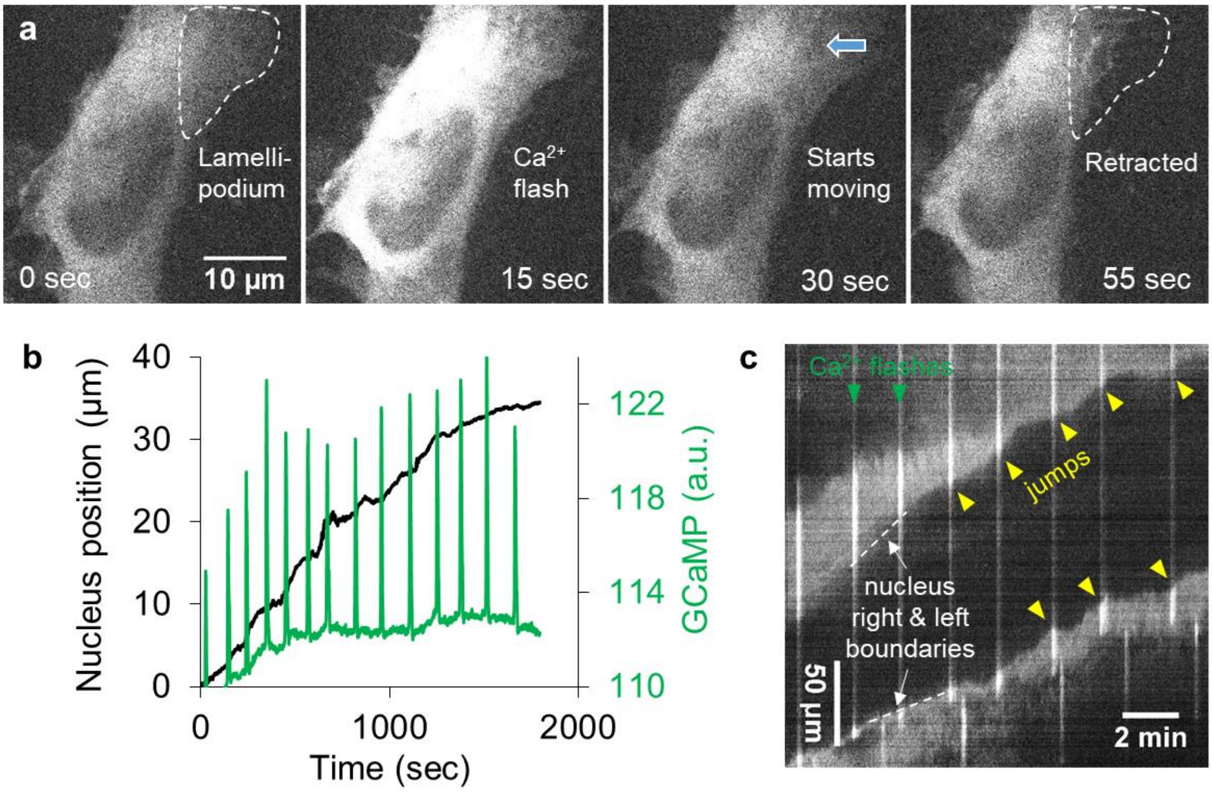


***Figure S17. Ca$^{2+}$ flashes induced discontinuities in ENCC migration on a 2D substrate.*** *(a) Stills from x60 timelapse of 2D ENCC culture showing lamellipodium retraction immediately after a Ca$^{2+}$ event (Video S10). (b) Nucleus centroid position of an ENCC migrating in 2D along the x axis (black). Ca$^{2+}$ transients (green) are correlated here 12 out of 14 times with cell speed discontinuities (Video S11). (c) Kymograph corresponding to the first 900 sec of (b) showing the nucleus right and left border and discontinuities (yellow arrowheads) associated with Ca$^{2+}$ events (white vertical lines).*
